# Induced Proteinuria Enhances Adeno-Associated Virus Transduction of Renal Tubule Epithelial Cells After Intravenous Administration

**DOI:** 10.1101/2025.05.28.656514

**Authors:** Jeffrey D. Rubin, Katayoun Ayasoufi, Erin B. McGlinch, Michael J. Hansen, Peter C. Harris, Vicente E. Torres, Michael A. Barry

**Author notes:** Corresponding author Correspondence to: Michael A. Barry, PhD Mayo Clinic 200 First Street SW Rochester, MN, USA.

## Abstract

A variety of genetic diseases of the kidney tubule are amenable to correction via gene therapy. However, gene delivery to renal tubule epithelial cells mediated by viral vectors via the blood is inefficient due to the permselectivity of the glomerular barrier. We hypothesized that effacement of podocyte foot processes would disrupt typical glomerular limitations on filtration and make renal tubule epithelial cells susceptible to transduction from viral vectors delivered intravenously. We determined that adeno-associated virus serotype 8 (AAV8) transduced significantly more epithelial cells in the kidney under the conditions of LPS-induced proteinuria. Use of AAV1 in tandem with LPS-induced proteinuria yielded an ideal two-pronged effect of both partially detargeting the liver and transducing the kidney with a higher bioluminescent signal than AAV8 did, and at half of the dose. Using adenovirus serotype 5 (Ad5) in conjunction with LPS-induced proteinuria showed that kidney transduction was enhanced, but only in glomerular cells. These studies mechanistically test the efficacy of different viral vectors and demonstrate their capacity to transduce kidney epithelial cells. This is a fundamental step in designing future treatments for kidney gene therapy.

**Significance Statement:** The slit diaphragms in the glomerulus of the kidney are too narrow to allow most solute from the blood to enter the nephron. As a result, large particles such as viral vectors generally cannot access tubule epithelial cells necessary to correct genetic disorders. In this study, mice are induced into a state of proteinuria and subsequently administered viral vectors intravenously. This group of mice had higher levels of transduced kidney epithelial cells than the control (non-proteinuria) group. The induction of transient proteinuria before intravenous viral vector administration is a novel approach that could make tubulopathies treatable via gene therapy in both animal models and humans.

## Introduction

While there have been many advances in the field of gene therapy in recent years, the kidney is one organ that has lagged behind others such as the eye, the nervous system, and the liver. Kidney diseases that affect the tubule of the nephron, i.e., tubulopathies, such as polycystic kidney disease, have yet to be successfully treated via gene therapy. The kidney is a particularly difficult target for viral vectors delivered intravenously. The slit diaphragm constitutes the physiological filtration system in the glomerular basement membrane of the nephron. These structures allow molecules of ≤10 nanometers (nm) and ≤50 kilodaltons (kDa) to pass from the blood to the proximal tubule of the nephron. Common viral vectors such as lentiviruses, adenoviruses (Ad), and adeno-associated viruses (AAV), are approximately 120 nm, 90 nm, and 25 nm in diameter, respectively, and thus too large to efficiently pass the glomerular filter and transduce tubule epithelial cells in the kidney (reviewed in ^1^). Past studies have shown that intravenous injections of Ad and AAV in mammals can efficiently transduce cells in the glomerulus, but not renal tubule epithelium^2–11^. Previous work demonstrated that one way to bypass the glomerular filter and transduce tubule epithelial cells is to perform direct kidney injections of vectors^12–15^. While these injection methods have shown some ability to transduce kidney tubule epithelium, the transduction pattern is typically stochastic and not necessarily consistent enough to genetically correct a tubulopathy.

On a cellular level, slit diaphragms are composed of foot processes of podocytes in the glomerulus. Kidneys with podocyte defects may leak proteins from the blood, such as albumin, into the tubule epithelium and eventually the urine in a condition known as proteinuria. We hypothesized that a state of induced proteinuria may allow the penetration of viral vectors further into the nephron where they would be able to transduce tubule epithelial cells. In 2004, Reiser *et al.* demonstrated that a state of induced proteinuria can effectively be achieved in mice via an intraperitoneal (i.p.) injection of lipopolysaccharide (LPS) through direct activation of toll-like receptor 4 present on podocytes rather than immune cell activation^16^. Thus, we proceeded to characterize the pharmacodynamic profile of gene therapy vectors during a state of induced proteinuria in mice.

Our primary goal was to determine if a state of proteinuria allowed any of these vectors to penetrate further into the nephron and transduce tubule epithelial cells compared to control. Using confocal microscopy, bioluminescence, and flow cytometry, we found that AAV8 significantly enhanced transduction of tubule epithelial cells in the kidney. The current study lays the groundwork for enhancing AAV-mediated gene delivery to renal tubule epithelium of mice and humans in a state of proteinuria or in which a state of induced proteinuria can be achieved.

## Materials and Methods

### Animal studies

Animals were housed in the Department of Comparative Medicine animal facilities at Mayo Clinic. All experiments and procedures were carried out according to the provisions of the Animal Welfare Act, PHS Animal Welfare Policy, the principles of the NIH Guide for the Care and Use of Laboratory Animals, and the policies and procedures of the Institutional Animal Care and Use Committee at Mayo Clinic. Both male and female mice were used.

### AAV vectors

AAV vectors were produced using a standard triple transfection and iodixanol gradient purification method^17^. Briefly, a vector plasmid (pTRS-CBh-Cre), a *rep* and *cap* plasmid (pRC), and a pHelper plasmid were transfected into 293T cells using polyethylenimine. Three days later, cells were harvested and lysed by successive freeze/thaw cycles. Cell lysate was overlayed onto an iodixanol gradient and ultracentrifuged for two hours. The banded AAV was extracted via needle and syringe and titrated via qPCR using SYBR™ Green. All AAV vectors used in this study were self-complementary (scAAV) with a cytomegalovirus chicken β-actin hybrid promoter (CBh) driving expression of the Cre recombinase gene.

### Ad vectors

Replication-defective Ad vectors were produced in 293 cells and purified by double banding on CsCl gradients. Cre expression is driven by the CMV promoter.

### Flow cytometry

Kidney samples were chopped into small pieces using scissors and put in Miltenyi© tubes. 2.35 mL of Gibco DMEM (cat # 11054001), 100 μL of enzyme D, 50 μL of enzyme R, and 12.5 μL of enzyme A from the Miltenyi “Tumor Dissociation Kit” were added into each sample. Samples were homogenized using soft tissue dissociation program on Miltenyi OctoMACS™ Separator. Samples were passed through 70 μM filters and spun at 400 xg for 10 minutes. Pellets were resuspended in 3.1 mL of cold DPBS, supplemented with 900 μL of Miltenyi Debris removal solution, overlayed with 4 mL of ice cold DPBS, and spun at 3000 g for 10 minutes. The samples were then washed with DPBS and red blood cells were lysed with 1 mL of ACK Lysis buffer for 1 minute. The samples were resuspended in 900 uL of RPMI and filtered using 35 μM flow tube filters.

Fluorescent staining occurred as follows: After all samples were processed and passed through filters, they were washed twice with PBS. Samples were stained with a master mix composed of conjugated antibodies against EpCAM PECy7 (1:250) [BioLegend, Cat# 118216], CD31 AF647 (1:500) [BioLegend, Cat# 102516], CD45 perCP (1:1000) [BioLegend, Cat# 103130], TCRβ BV421 (1:1000) [BioLegend, Cat# 109229], CD4 BV510 32 μL (1:500) [BioLegend, Cat# 100449], CD8a BV570 (1:500) [BioLegend, Cat# 100739], CD11b BV650 (1:1000) [BioLegend, Cat# 101259], Ghost Dye Red 780 (1:2000) [Tonbo Biosciences, Cat# 13-0865-T100], and FC block (1:500) [BD Pharmingen, Cat# 553141]. Three minutes prior to experimental mice being sacrificed, 3 μg of CD45 BV711 [BioLegend, Cat #103147] was injected intravenously to be able to distinguish between circulating and tissue resident CD45+ cells. Samples were stained for 30 minutes at 4C in the dark, washed twice with PBS and ran on Cytek™ Aurora spectral 4-laser flow cytometer.

For the experiments also staining against α-Fucose, Lotus Tetragonolobus Lectin (LTL), Biotinylated (1:100) [Vector Laboratories, Cat# B-1325-2] was the primary stain and BV786 Streptavidin (1:2000) [BD Horizon, Cat# 563858] was the secondary stain. For the experiments also staining against Aquaporin-1, Anti-Aquaporin-1 (1:100) [Boster Biological Technology, Cat# PB9473] was the primary stain and anti-rabbit AF647 (1:2000) [Invitrogen, Cat# A-21245] was the secondary stain. In these experiments, CD31 was stained using anti-CD31 BV510 (1:150) [BD Biosciences, Cat# 740124].

### In vivo bioluminescent imaging

Mice were anesthetized with isoflurane and injected intraperitoneally with 150 μL of D-Luciferin (20 mg/ml; RR Labs Inc., San Diego, CA).

Images were taken using PerkinElmer IVIS® Lumina S5 Imaging System ten minutes after D-Luciferin administration and luminescence was quantified using Living Image software. During *ex vivo* tissue imaging, tissues were placed in either 6-well or 12-well tissue culture vessels and imaged. In all cases except for Figure 1, kidneys were laterally bisected before imaging to prevent squelching of luminescence by the kidney capsule.

**Figure 1.**
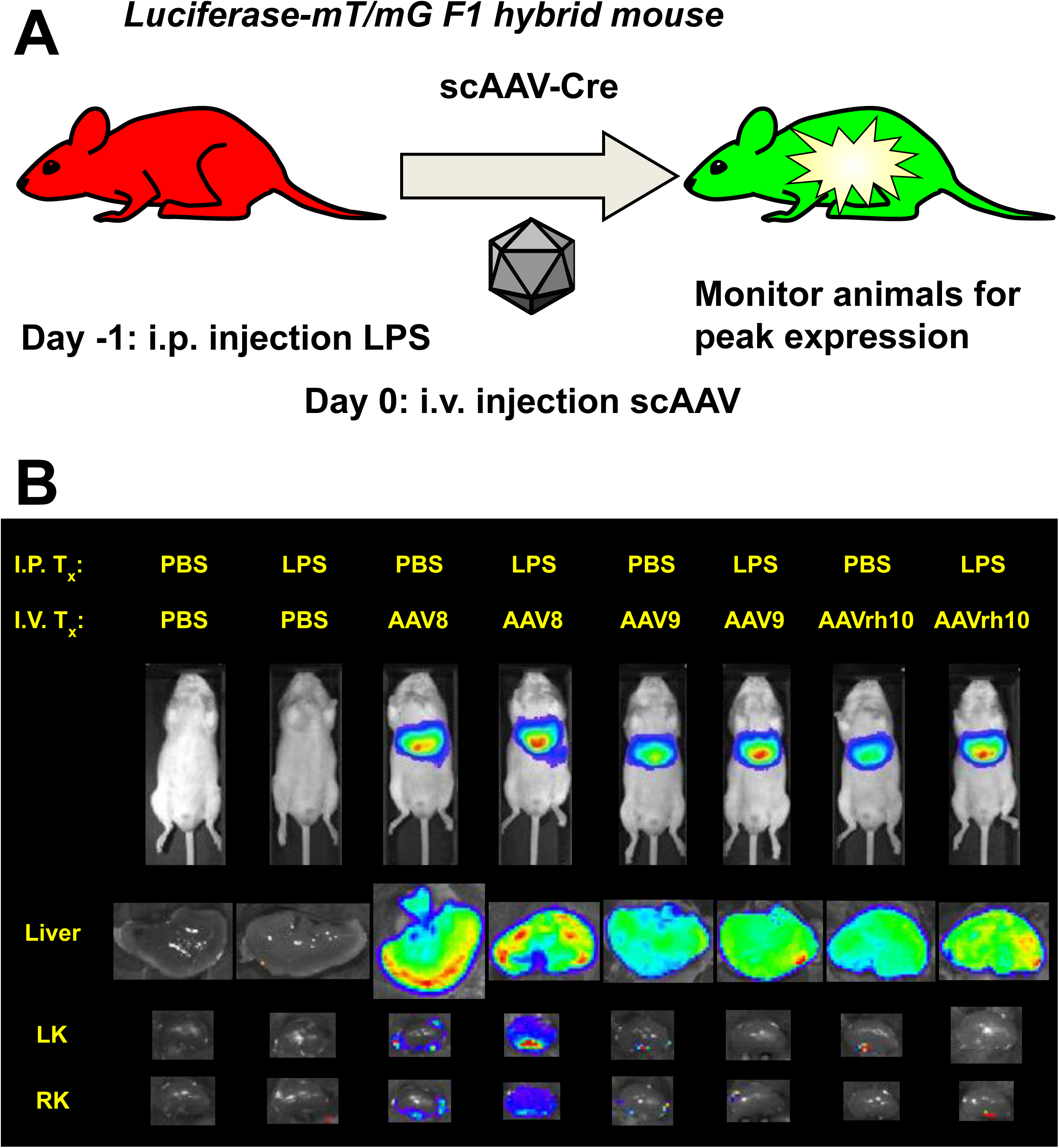
Intravenous delivery of AAV8 in a state of induced proteinuria enhances kidney transduction. **(A)** Diagram of experimental scheme. Two month old male luciferase-mT/mG triple reporter mice were administered LPS intraperitoneally on Day -1 and scAAV intravenously on Day 0. *In vivo* bioluminescence was assessed daily until peak expression was observed at Day 6. **(B)** *In vivo* bioluminescence at Day 6 followed by *ex vivo* luminescence of livers and kidneys. *n* = 1 mouse per group.

### Statistical Analyses

All statistical analyses were performed using GraphPad Prism 9. All p-values were generated using Mann-Whitney tests unless otherwise noted. All tests were two-tailed. Error bars and exact *n* values used to calculate statistics are described in corresponding figure legends.

### Tissue sectioning and confocal microscopy

Tissues from mice with membrane-bound fluorescent proteins were fixed by overnight immersion in 4% paraformaldehyde (PFA)-PBS at 4°C, then cryoprotected overnight in 15% sucrose-PBS and 30% sucrose-PBS, successively, at 4°C. Trimmed tissues were then flash frozen by dry ice-cooled isopentane in optimal cutting temperature (O.C.T.) medium (Sakura Finetek). Cryosections (18 μm thickness) were prepared with a Leica CM1860 UV cryostat (Leica Biosystems) and mounted on slides (Superfrost Plus; Thermo Fisher Scientific, Waltham, MA) with VECTASHIELD with 4’,6-diamidino-2-phenylindole (DAPI) (Vector Laboratories, Burlingame, CA), and CytoSeal-60 coverslip sealant (Thermo Fisher Scientific). Confocal imaging was performed at the Microscopy and Cell Analysis Core facility at Mayo Clinic Rochester (Rochester, MN), using a Zeiss LSM780 laser confocal microscope (Carl Zeiss Jena, Jena, Germany).

For tissue sections stained with lotus tetragonolobus lectin (LTL), the slides containing tissue sections were washed with PBS, treated with 5% normal goat serum (Abcam Catalog # ab7481) and 0.5% IGEPAL® CA-630 (Sigma I8896) dissolved in PBS blocking buffer for 1 hour at room temperature. The slides were then incubated with a 1:100 dilution of biotinylated LTL (Vector Laboratories Cat. No: B-1325) overnight at 4°C. The slides were washed and then incubated with a 1:200 dilution of streptavidin-Alexa Fluor 647 (Invitrogen Catalog # S21374) at room temperature for one hour. The slides were washed and coverslips were mounted using Vectashield (without DAPI).

### Transgenic mice

LSL-Luc mice (Stock No: 005125) and mT/mG mice (Stock No: 007576) were originally purchased from The Jackson Laboratory.

## Results

### AAV8 gene delivery to the kidney is increased in a state of induced proteinuria

To investigate the effects of induced proteinuria on viral vector gene delivery to the kidney, we utilized LPS. Mice were administered an i.p. injection of 200 μg of LPS, the mode of delivery and dose concluded to be effective in previous work^16^. The following morning, urine was collected from mice injected with either LPS or PBS as a control and assayed using a proteinuria dipstick to ascertain whether proteinuria had effectively been induced (**Supplemental Figure 1**). Subsequently, mice were administered i.v. injections of self-complementary AAV8-Cre (scAAV8-Cre), scAAV9-Cre, scAAVrh10-Cre, or PBS as control (*n* = 1 for each combination of PBS or LPS and each vector). The mice used in this experiment were LSL-Luc-mT/mG F1 hybrid mice: each mouse has one LoxP-STOP-LoxP-Luciferase allele and one tdTomato/EGFP (mT/mG) allele at the ROSA locus. Thus, each mouse has Luciferase and mG genes activatable by Cre-expressing vectors, allowing for tracking of vector pharmacodynamics on both a cellular and tissue-specific level (**Figure 1A**)^10,15^.

Bioluminescent imaging was performed until the signals reached an approximate plateau at day 6 (**Supplemental Figure 2A**). Due to the high liver tropism of the three AAV serotypes used, the signals measured *in vivo* were almost certainly emitted from the livers of these mice (**Figure 1B**). To directly assess liver and kidney transduction, the mice were sacrificed, and these organs were imaged *ex vivo*. While kidneys of the AAV9 and AAVrh10 injected mice exhibited minimal luminescence which was localized to the renal pelvis region of the kidney, the kidneys of the mouse with induced proteinuria injected with AAV8 had pervasive luciferase expression throughout the entire kidney. This observation indicates a clear difference in vector pharmacodynamics in mice in states of induced proteinuria.

To assess kidney transduction on a cell-by-cell basis, we examined direct fluorescence via confocal microscopy. We observed that treating mice with LPS prior to AAV injection resulted in many instances of transduced cells with tubular morphology adjacent to glomeruli (**Figure 2**). We hypothesized that after podocyte foot process effacement, viral vectors might filter through the glomerulus and contact the most proximal part of the nephron, the proximal tubule. We counterstained kidney sections with lotus tetragonolobus lectin (LTL), a marker of proximal tubule cells. Surprisingly, no instances of EGFP^+^ transduced cells seemed to be double positive for the LTL stain.

**Figure 2.**
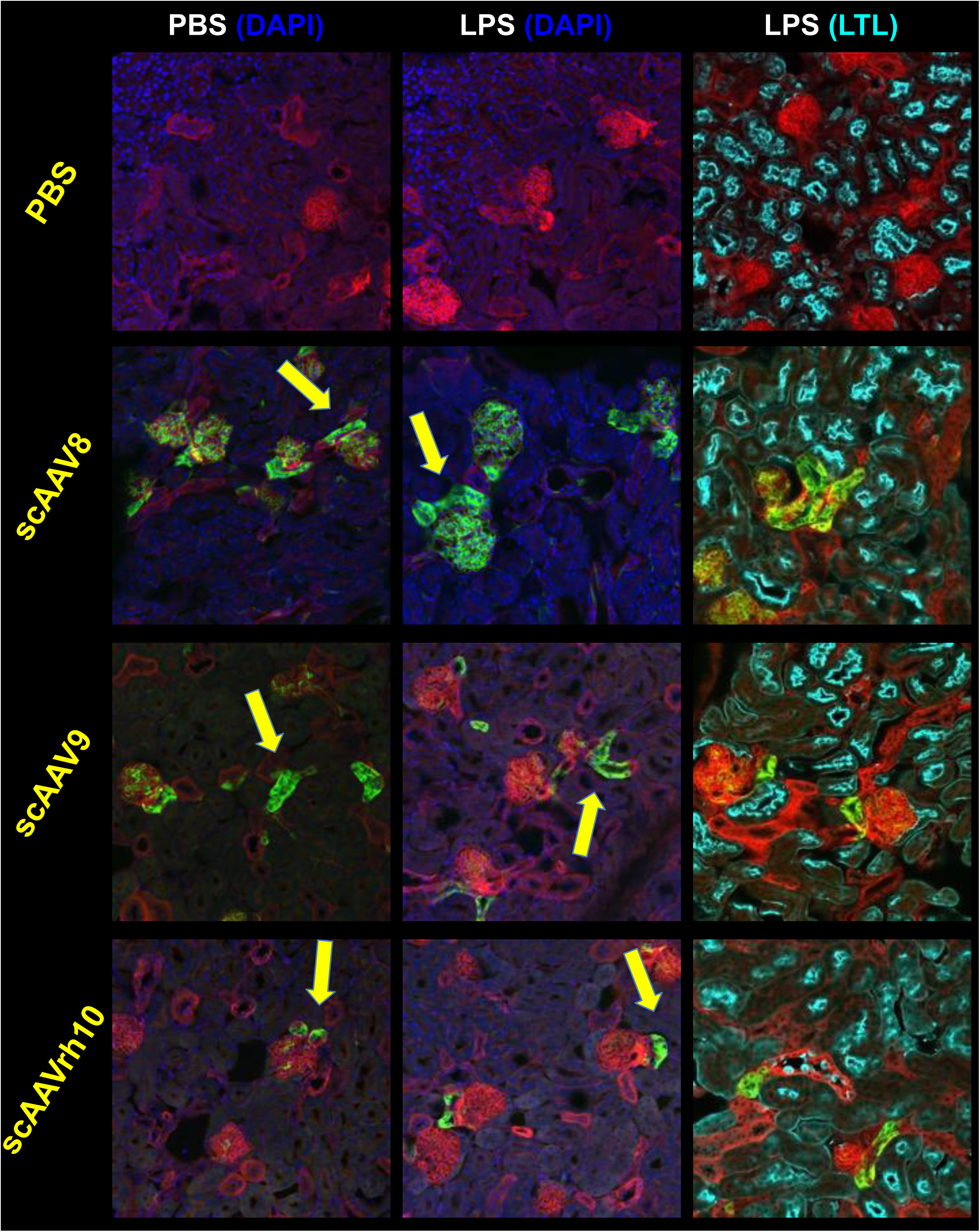
Intravenous delivery of multiple AAV serotypes enhances tubule epithelial cell transduction, but not necessarily proximal tubule cell transduction. The same kidneys from Figure 1 were sectioned to examine endogenous mT and mG fluorescence. Arrows point to examples of transduced non-glomerular (tubular) cells. While some tubular cell transduction was observed in PBS-injected control mice (left panels), there were increased numbers of these cells in LPS-injected induced proteinuria mice (center panels). No instances of these transduced cells were observed to be counterstained by LTL, a marker of proximal tubule cells (right panels). *n* = 1 mouse per group.

This indicates that although induced proteinuria allows AAV to penetrate further into kidney tissue from the blood and transduce more tubule cells, these cells are not necessarily proximal tubule cells.

### AAV8 significantly increases renal epithelial cell transduction during proteinuria

AAV serotypes 8, 9, and rh10 each potentially increased transduction of renal tubule epithelial cells in an induced state of proteinuria, with AAV8 having the most striking effect (**Figure 1B**). To quantify this effect, new groups of mice were given an i.p. administration of either PBS or LPS at Day -1 and an i.v. administration of scAAV8-Cre at Day 0 at a dose of 2e11 genome copies (GC). Proteinuria dipsticks from these groups of mice at Day -1 (baseline) and Day 0 (post PBS or LPS) are shown as examples (**Supplemental Figure 1**).

Mice were imaged for *in vivo* luminescence at Day 6 at which point the mice were sacrificed and their tissues imaged *ex vivo*. There was no significant difference observed between PBS and LPS-injected groups *in vivo* or liver *ex vivo* (**Figure 3A**).

**Figure 3.**
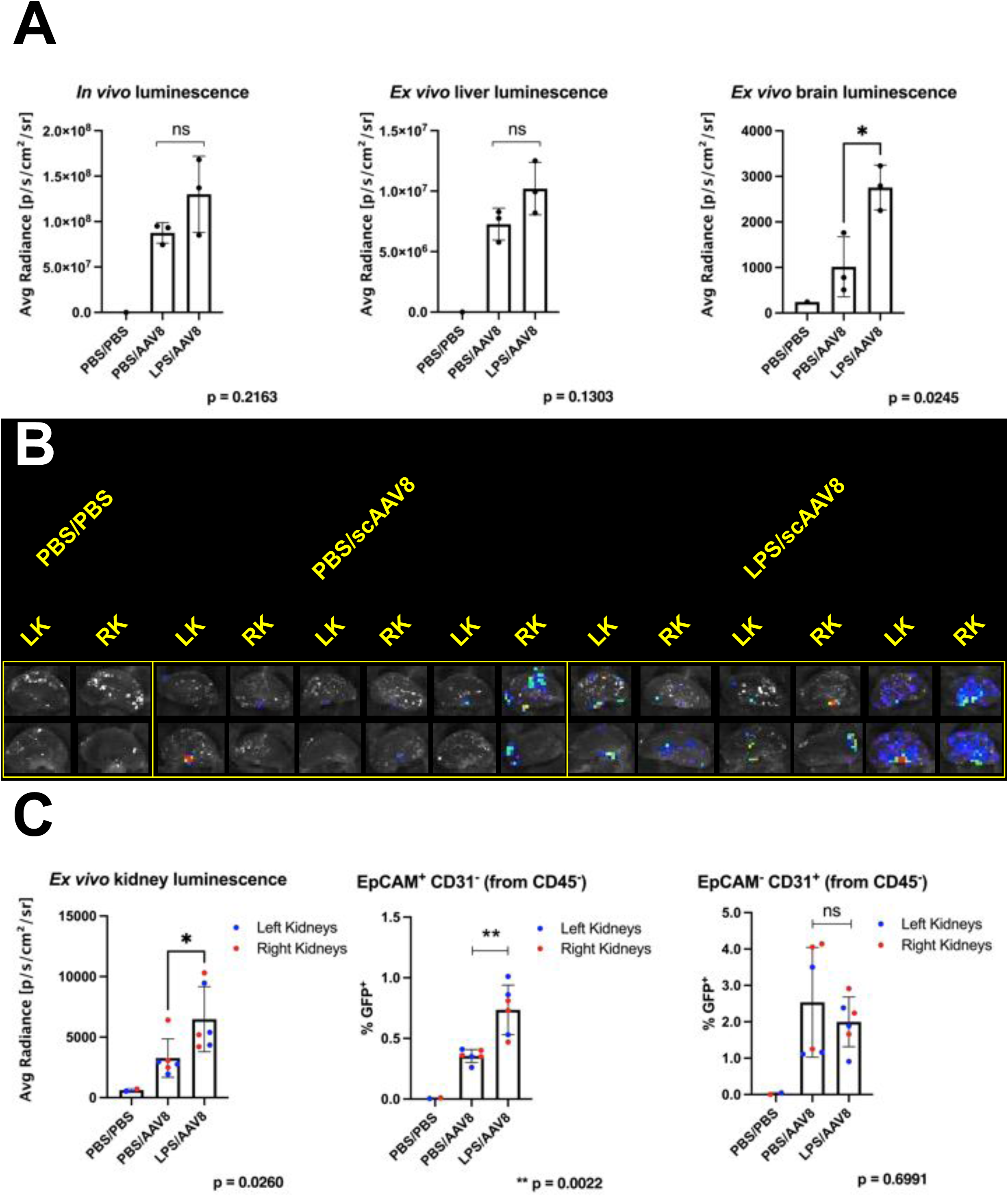
Intravenous delivery of scAAV8 in a state of induced proteinuria significantly increases transduction of renal epithelial cells. **(A)** Three month old male mice were administered an i.p. injection of either PBS or LPS at Day -1 and an i.v. injection of 2.03e11 GC of scAAV8-Cre at Day 0. At Day 6, *in vivo* luminescence and *ex vivo* liver luminescence were not significantly different between PBS and LPS-injected groups, although brain luminescence was significantly increased in the LPS-injected group (p = 0.0475 by Welch’s t test.). *n* = 3 mice per group, except for control group where *n* = 1; error bars are represented by mean with SD. **(B)** Kidneys were bisected with a razor blade to reduce obstruction of luminescence and imaged *ex vivo*, with the LPS-injected group exhibiting increased luminescence compared to the PBS-injected group. (C) *Ex vivo* luminescence from Panel B was quantified and kidneys were subsequently processed for flow cytometry. Overall, kidneys from LPS-injected mice showed significantly higher *ex vivo* luminescence and percentage of transduced epithelial cells, but not of transduced endothelial cells (p values obtained using Mann-Whitney test). *n* = 6 kidneys per group, except for control group where *n* = 1; error bars are represented by mean with SD.

Brain luminescence was significantly increased, indicating that LPS administration induced some blood brain barrier disruption^18–20^ prior to AAV injection. In contrast to the livers, *ex vivo* kidney transduction visibly increased in LPS-injected mice versus PBS-injected mice (**Figure 3B**). The kidneys of the LPS-injected mice exhibited significantly higher luminescence than those of the control mice (**Figure 3C**). These kidneys were then processed for flow cytometry and labeled to detect epithelial cell adhesion molecule (EpCAM), a marker of epithelial cells, and CD31, a marker of endothelial cells. It was found that epithelial cells, but not endothelial cells, had a significant increase in transduction, indicating that the injected AAV8 did in fact have more access to epithelial cells during a state of induced proteinuria. (**Figure 3C**). In addition, a significantly increased quantity of EGFP^+^ macrophages were found in the blood of LPS-injected mice as compared to PBS-injected mice, indicating that an increased presence of macrophages was induced by LPS administration and subsequently transduced by scAAV8-Cre (**Supplemental Figure 3B**). Representative flow plots and gating strategies are shown in **Supplemental Figure 4**.

### AAVrh10 does not increase epithelial cell transduction during proteinuria

We next sought to determine if a particular serotype of AAV could in fact result in a significantly increased number of epithelial cells in the kidney during induced proteinuria. Initially, AAV8 had stronger results than AAV9 or AAVrh10 (although this variation could have been due to a difference in dosing). To quantify the effects of a serotype other than AAV8, we repeated the prior flow cytometry experiment using scAAVrh10-Cre rather than scAAV8-Cre. We first examined the %EGFP^+^ (transduced) cells present among CD45^-^ (non-hematopoietic) and CD45^+^ (hematopoietic) cells in the kidneys (**Figure 4A**). Interestingly, there was no difference in transduced CD45^-^ cells between PBS and LPS-injected groups. However, there was a significant increase in transduced CD45^+^ cells. This effect indicates that after LPS administration, either more CD45^+^ cells mobilized in the blood or an increased number of hematopoietic cells infiltrated the kidney and had increased susceptibility to transduction.

**Figure 4.**
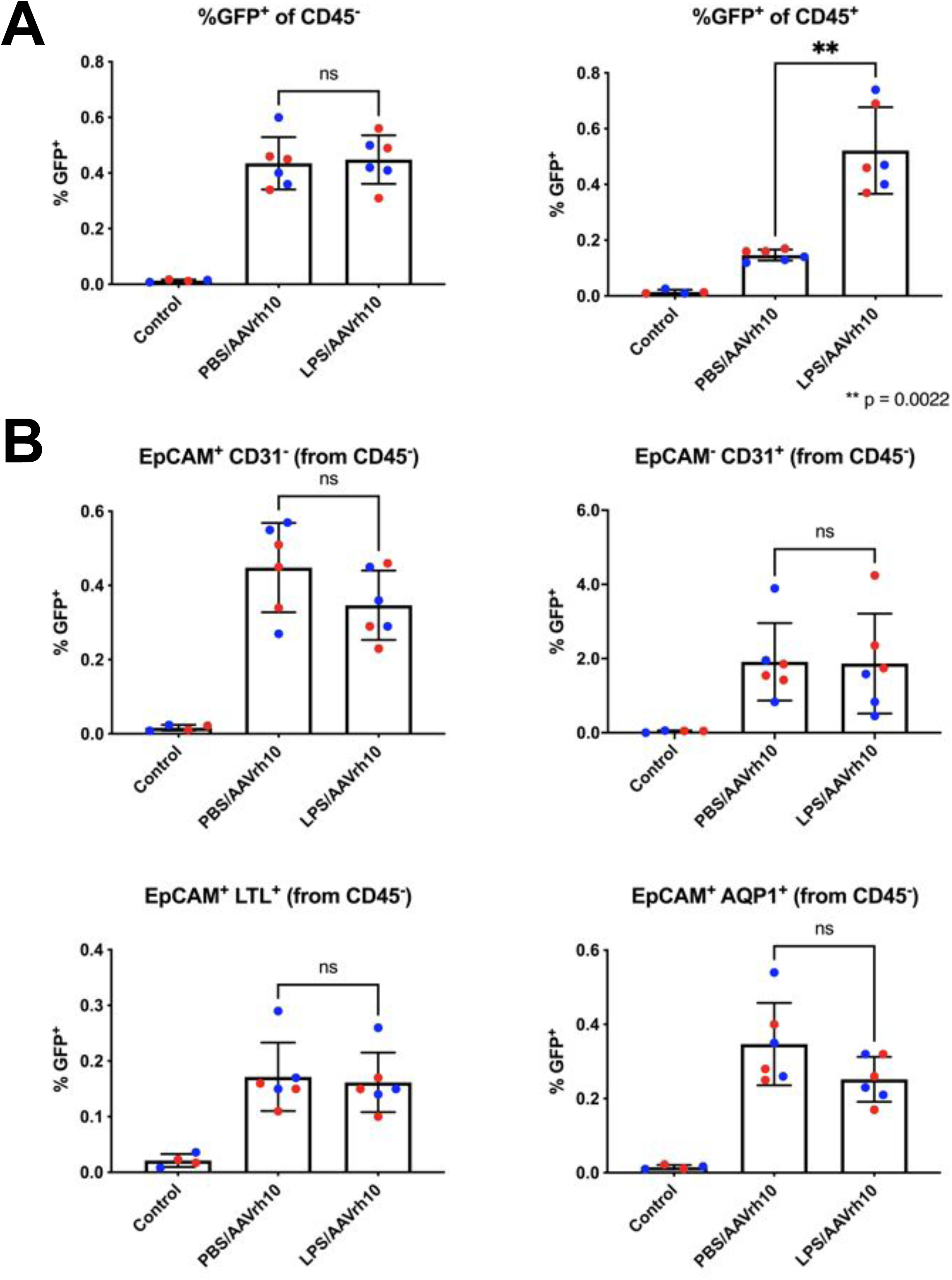
AAVrh10 does not necessarily increase transduction of tubule epithelial cells during induced proteinuria. **(A)** Eight month old female mice were administered an i.p. injection of either PBS or LPS at Day -1 and an i.v. injection of 1.76e11 GC of scAAVrh10-Cre at Day 0. At Day 5, kidneys were processed for flow cytometry. Although there was no difference in transduced CD45^-^ (non-hematopoietic) kidney cells, kidneys of the LPS-injected group had a significant increase in CD45^+^ (hematopoietic) cells compared to the PBS-injected group (*n* = 6 kidneys per group). **(B)** Kidney CD45^-^ (non-hematopoietic) cells were separately gated into EpCAM^+^ CD31^-^ (all epithelial cells), EpCAM^-^ CD31^+^ (endothelial cells), and EpCAM^+^ LTL^+^ and EpCAM^+^ AQP1^+^ (two different markers of proximal tubule cells). None of the aforementioned gating strategies showed a significant difference in transduced cells between PBS-injected and LPS-injected groups. *n* = 6 kidneys per group, except for control group where *n* = 1; error bars are represented by mean with SD for all panels. p values determined using Mann-Whitney tests for all panels.

We next evaluated the transduction of epithelial cells in the kidney. We first examined the %EGFP^+^ cells amongst CD45^-^ EpCAM^+^ and CD45^-^ CD31^+^ populations, which represent transduced epithelial cells and transduced endothelial cells, respectively. When this experiment was performed using scAAV8 (**Figure 3**), there was a significant increase in transduced epithelial cells, but not endothelial cells, between the LPS and PBS-injected mice. However, when this experiment was repeated using scAAVrh10, there was no significant difference between the LPS and PBS-injected groups of mice (**Figure 4B**). To further clarify the extent to which proximal tubule cells were transduced, samples were also labeled with LTL and aquaporin-1 (AQP1). In both cases, no significant difference was observed between transduced cells in LPS or PBS-injected mice. Overall, mice with or without induced proteinuria did not seem to have a difference in transduced renal epithelial cells after intravenous injection of scAAVrh10-Cre. Representative flow plots and gating strategies are shown in **Supplemental Figure 5**.

### A vector with low liver tropism enhances kidney transduction during induced proteinuria

Although increasing transduction in tubule cells in the kidney is an important goal for efficacy of gene therapy, detargeting vectors from off-target tissues is an important facet of gene therapy safety^21,22^. While AAV8 showed efficacy in terms of increasing kidney transduction during a state of induced proteinuria, it also fully transduces the liver (**Supplemental Figure 3A**). To attempt to resolve the off-target tissue transduction, we tested AAV1, a serotype known to have lower liver tropism than other serotypes, in conjunction with induced proteinuria^23^.

Mice were administered an i.p. injection of either PBS or LPS at Day -1 and an i.v. injection of scAAV1-Cre at Day 0 at a dose of 9.5e10 GC. Similar to previous experiments, *in vivo* luminescence signals peaked at Day 6, at which point mice were sacrificed and *ex vivo* liver luminescence was comparable between both groups of mice (**Figure 5A**, top). Although mean kidney *ex vivo* luminescence was increased in LPS-injected mice versus PBS-injected mice, the difference was not significant. Interestingly, when comparing this data side-by-side with the *ex vivo* kidney luminescence data of the scAAV8-Cre injected mice from **Figure 3**, both PBS-injected and LPS-injected groups of scAAV1-injected mice had higher signals than the LPS and scAAV8-injected mice, and at approximately half of the dose of scAAV8, indicating that scAAV1 may have a higher native kidney tropism both with and without induced proteinuria (**Figure 5A**, lower). Histologically, injected with PBS followed by scAAV1 had many instances of transduced glomerular cells while mice injected with LPS followed by scAAV1 had increased instances of transduced tubular cells (**Figure 5B**). Importantly, the livers of the mice injected with scAAV1 were only partially transduced, while the livers of the mice injected with scAAV8 were fully transduced (**Supplemental Figure 7**). These data indicate that scAAV1 may be an ideal vector for targeting renal tubule epithelial cells while avoiding unnecessary transduction of hepatocytes.

**Figure 5.**
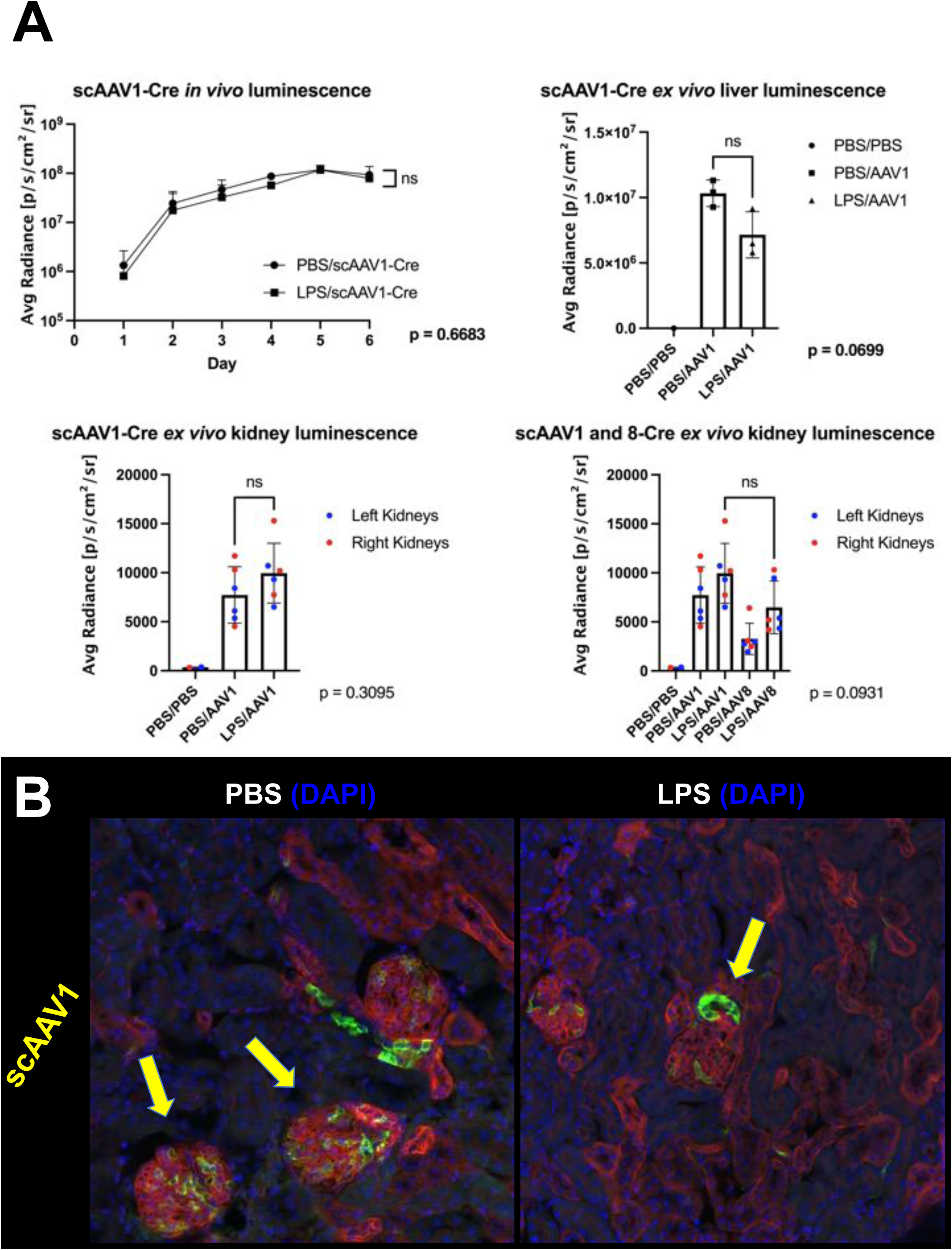
Examination of kidney transduction using a vector with low liver tropism. **(A)** Four and a half month old female mice were administered an i.p. injection of PBS or LPS on Day -1 and an i.v. injection of 9.5e10 GC of scAAV1-Cre on Day 0. *In vivo* bioluminescence was assessed daily until peak expression was observed at Day 6. No significant difference was observed between groups, including a measurement of *ex vivo* liver luminescence (p values determined using Welch’s t test). *n* = 3 mice per group, except for control group where *n* = 1. Error bars are represented by mean with error (top left) or mean with SD (top right). *Ex vivo* kidney luminescence showed that LPS-injected mice had an increased but insignificant amount of luminescence compared to PBS-injected mice as well as LPS and scAAV8-Cre injected mice (p values determined using Mann-Whitney test). *n* = 6 kidneys per group, lower panels; error bars are represented by mean with SD; scAAV8-Cre data represents the same data shown in Figure 3. **(B)** The kidneys analyzed in Panel A were sectioned to observe endogenous mT and mG fluorescence. While mice treated with PBS and scAAV1-Cre showed transduction primarily in glomeruli (left), mice treated with LPS and scAAV1-Cre showed increased transduction in non-glomerular (tubular) cells (right). Arrows point to examples of transduced glomerular cells (left) or examples of transduced tubule cells (right).

### Induced proteinuria enhances Ad5 transduction of glomerular, but not epithelial cells

Notable differences in the transduction profiles of kidney and liver cells were observed in the various AAV capsids tested. The variation in transduction profiles is likely due to differences in receptor usage as well as capsid surface electromagnetic charges^24–26^. To test other applications and potential limitations of the induced proteinuria method, we next used a physically larger gene delivery vector, replication-defective adenovirus serotype 5 expressing Cre recombinase (Ad5-Cre).

Mice were administered i.p. injections of either PBS or LPS on Day -1 and i.v. injections of Ad5-Cre on Day 0. *In vivo* luminescent signals (indicative of liver transduction) were monitored up to Day 5 until they peaked. In contrast to previous experiments using AAV, mice injected with LPS prior to Ad5-Cre had significantly reduced *in vivo* luminescence compared to PBS-injected mice (**Figure 6A**, left). The mice were then sacrificed and their kidneys were imaged for *ex vivo* luminescence. Kidneys from mice administered LPS prior to Ad5-Cre had a significantly higher signal than those from the control group (**Figure 6A**, right). In the images of the kidneys of PBS and Ad5-Cre injected mice, little to no luminescence is visible, whereas in the kidneys of the LPS and Ad5-Cre injected mice, two out of three of the kidneys showed enhanced luminescence localized near the renal pelvis (**Figure 6B**), in contrast to kidney images of mice injected with LPS and scAAV8, which showed more diffuse luminescence throughout the kidney (**Figure 3**).

**Figure 6.**
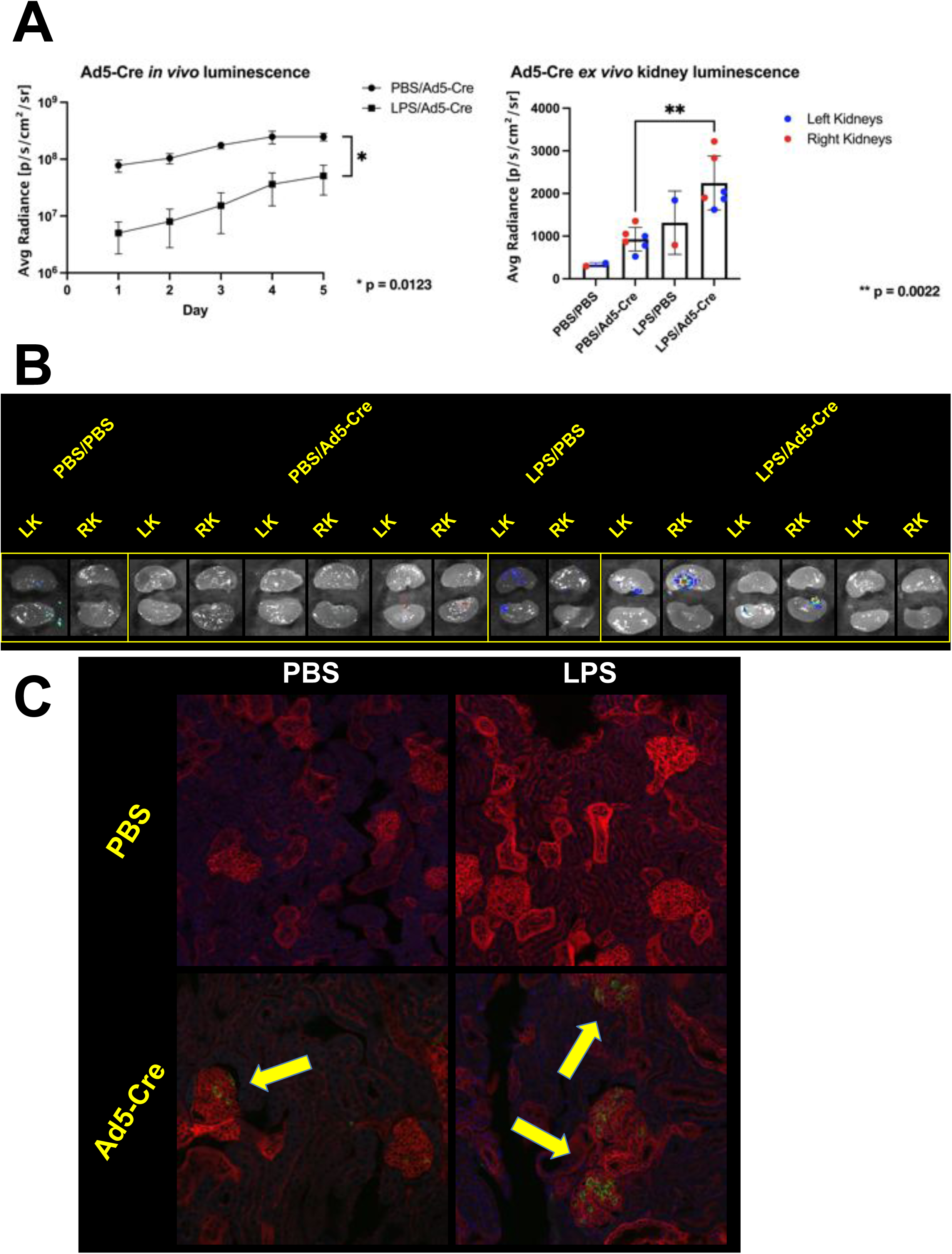
Induced proteinuria increases adenovirus transduction of the kidney, but strictly in glomeruli. **(A)** Four month old mice were administered an i.p. injection of PBS (male mice) or LPS (female mice) on Day -1 and an i.v. injection of 1e11 vp of Ad5-Cre on Day 0. *In vivo* bioluminescence was assessed daily until peak expression was observed at Day 5. Luminescence was significantly lower in LPS-injected mice compared to PBS-injected mice (p value determined using Welch’s t test; *n* = 3 mice per group; error bars are represented by mean with error, left), however, *ex vivo* kidney luminescence was significantly higher in LPS-injected mice compared to PBS-injected mice (p value determined using Mann-Whitney test; *n* = 6 kidneys per group, except for control group where *n* = 1; error bars are represented by mean with SD, right). **(B)** Bioluminescent images of the kidneys quantified *ex vivo* in Panel A. While essentially no luminescence is visible in kidneys from mice injected with PBS, kidneys from mice injected with LPS showed luminescence localized to the renal pelvis region of the kidney. **(C)** Kidneys shown in Panel B were sectioned to examine mT and mG endogenous fluorescence. Yellow arrows point to examples to transduced glomerular cells, which are present sparsely in mice injected with PBS and more frequently in mice injected with LPS. No instances of transduced tubular cells were observed in either group of mice.

Notably, in contrast to previous experiments using AAV, no instances of transduced tubule cells were observed in kidneys of PBS or LPS and Ad5-Cre injected mice. However, there were observed to be an increased number of glomerular cells transduced in the LPS and Ad5-Cre injected mice versus the PBS-injected mice, indicating that induced proteinuria did not enhance penetration of Ad5-Cre into renal epithelial tubular cells but may have aided penetration further into the glomerulus itself (**Figure 6C**). Liver histology showed that mice injected with PBS followed by Ad5-Cre had fully transduced livers while mice injected with LPS followed by Ad5-Cre had only partially transduced livers, possibly as a result of LPS interactions with Kupffer cells (**Supplemental Figure 6A**). These data indicate that while the induced proteinuria method used in tandem with Ad may not necessarily be effective in treating tubulopathies, it may be helpful in increasing gene delivery to glomerular cells.

## Discussion

Previous work has strived to pinpoint on a tissue-to-tissue and cell-to-cell basis the tropisms of the various serotypes of AAV available for preclinical research in gene therapy^10,11,23^. These important studies increased knowledge of which capsids are appropriate for treatment of particular diseases based on transduction patterns.

However, these studies also highlighted the disappointingly low level at which renal tubule epithelial cells are transduced after systemic administration of AAV. Subsequent studies therefore focused on improving gene delivery to the kidney by finding ways to bypass the glomerulus, which is modeled as the stopping point of AAV and larger vectors, such as Ad, from the blood. These methods of direct kidney injection include renal arty infusion, renal vein injection, retrograde infusion into the ureter, and subcapsular injection, each of which has showed efficacy in tubule cell delivery over i.v. injection^12–15,27^. However, each of these techniques is invasive and results in stochastic cell transduction.

In the current study, we hypothesized that transiently effacing the podocyte processes in the glomerulus would allow increased penetration of viral vectors into the proximal tubule of the nephron. We showed through *ex vivo* bioluminescence that LPS-induced proteinuria increased overall transduction of the kidney after i.v. injection of several serotypes of AAV and a single serotype of Ad (**Figure 1**, **Figure 3**, **Figure 5A**, and **Figure 6**; results are summarized in **Table 1**). Upon sectioning these kidneys, we showed direct evidence of increased renal tubule transduction (**Figure 2** and **Figure 5B**). In addition, we showed that the percentage of kidney epithelial cells transduced after LPS-induced proteinuria significantly increased by flow cytometry (**Figure 3C**).

**Table 1.** Summary of kidney transduction profile of five different gene therapy vectors with and without induced proteinuria.

| Vector | Known Receptors | Kidney Transduction (Baseline) | Kidney Transduction (Induced Proteinuria) |
| --- | --- | --- | --- |
| scAAV1 | α2,3 and α2,6 N-linked sialic acids, AAVR | Mostly glomerular | Increased tubular |
| scAAV8 | Laminin, AAVR | Mostly glomerular | Increased tubular |
| scAAV9 | Galactose, laminin, AAVR | Mostly glomerular | Increased tubular |
| scAAVrh10 | Unknown | Mostly glomerular | Increased tubular |
| Ad5 | CAR, integrin | Glomerular | Increased<br>glomerular |

However, we were unable to identify the additional transduced cells in the kidney as proximal tubule cells. Due to these cells’ consistent juxtaposition to glomeruli, they are likely to be cells of the macula densa, a section of the distal tubule that senses sodium chloride concentration and initiates release of renin^28^.

The current study shed light on several other aspects of kidney gene therapy. Firstly, using AAV8 in tandem with LPS-induced proteinuria resulted in a significantly increased quantity of transduced renal epithelial cells, while using AAVrh10 did not (**Figure 3** and **Figure 4**, respectively). This indicates vector tropism can be a prerequisite for AAV to transduce renal tubule epithelial cells. AAV8 uses laminin and AAVR as receptors but the receptors for AAVrh10 are unknown^25,29^. The AAV8 experiment was performed using male mice and the AAVrh10 experiment was performed using female mice. Seminal studies have shown that AAV is biased for increased transduction in tissues such as liver and heart in male versus female mice^30,31^. In addition, there may be unanticipated sex differences in kidney physiology that affect viral vector transduction, such as the recent discovery that macula densa cells project processes that reach toward the glomerulus and are longer in female mice than male mice^28^. Secondly, there was a significant increase in the quantity of transduced hematopoietic lineage cells (marked by CD45) in LPS-induced proteinuria mice versus control mice (**Figure 4A** and **Supplemental Figure 3B**). This effect seems most likely to occur due to a broad activation of immune cells at Day -1 when the i.p. injection LPS is administered. Perhaps the increased number of CD45^+^ immune cells in circulation, such as macrophages, accounts for the overall increase of transduction by AAV when administered at Day 0. Thirdly, AAV1 was not only effective as a liver detargeted vector but also as a renal-targeted vector. Mice without induced proteinuria administered an i.v. injection of AAV1 had higher *ex vivo* kidney luminescence than mice with LPS-induced proteinuria administered an i.v. injection of AAV8 (**Figure 5A**).

Strikingly, the aforementioned result was achieved when the dose of AAV1 was half the dose of AAV8 (9.5e10 GC versus 2e11 GC). The cellular receptors for AAV1 are α2,3-linked and α2,6-linked sialic acids, which are expressed at least in tubule epithelial cells distal to the loop of Henle^24,32^. Unnecessary, massive transduction of the liver can precipitate harmful side effects as evidenced by the death of Jesse Gelsinger in 1999 in a clinical trial to treat ornithine transcarbamylase deficiency and the more recent deaths of three patients in 2020 in a clinical trial to treat X-linked myotubular myopathy^33,34^.

Therefore, striking a balance by employing vectors reasonable kidney tropism and low liver tropism may be ideal.

Lastly, testing Ad5 made apparent some limitations of the LPS-induced proteinuria method. Induced proteinuria increased Ad5 transduction in the kidney compared to control, but only in the glomerulus (**Figure 6**). This data indicates that glomerular effacement had an effect on Ad pharmacodynamics in the kidney, but some segment of the glomerular structure still prevented Ad from penetrating into the tubular lumen. This is most likely due to the distinct size differences of AAV and Ad vectors, with the former being approximately 25 nm in diameter and the latter being approximately 90 nm in diameter (**Figure 7**). On a quantitative basis, the mean levels of *ex vivo* kidney luminescence attained by mice with induced proteinuria injected with scAAV1-Cre, scAAV8-Cre, and Ad5-Cre were 9962, 6480, and 2247, respectively (**Figure 5** and **Figure 6**). This data has implications for other large gene therapy vectors, such as lentiviruses, which are up to 120 nm in diameters, and lipid nanoparticle (LNP) complexes, which may have volumes beyond that of AAV. In terms of systemic effects of LPS, the extent to which LPS administration affected a particular mouse seemed to be generally correlated to lower liver transduction and higher kidney transduction when used with Ad5 (**Supplemental Figure 6B**).

**Figure 7.**
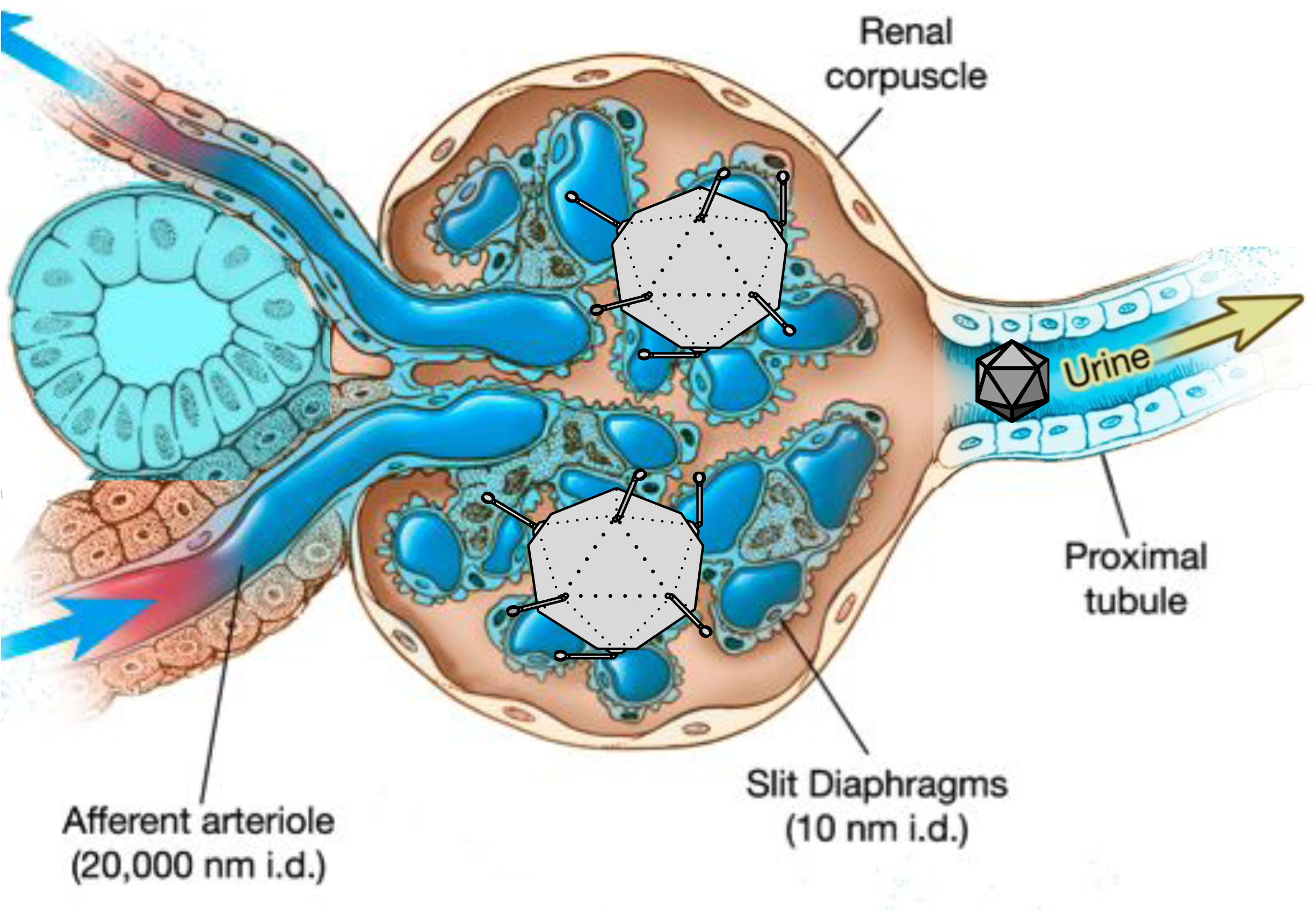
Diagram modeling vector pharmacokinetics in a state of induced proteinuria. LPS administration results in degradation of podocyte foot processes, effectively increasing the permselectivity of slit diaphragms to an unknown diameter above the natural 10 nm. This change in physiology allows the smaller AAV (25 nm i.d.) to penetrate into adjacent tubule cells while the larger Ad (90 nm i.d.) has increased penetration into glomerular cells but not tubular cells. It is also possible that AAV moved from the vasculature of the kidney to transduce cells of the macula densa.

Overall, the current study presents a novel method for consistently increasing transduction of renal tubule epithelial cells after i.v. injection of AAV. This method is advantageous due to its ability to consistently increase AAV delivery to tubule cells in both kidneys after a single intravenous injection. Although safer substances such as protamine sulfate have been shown in the literature to induce podocyte foot process effacement after renal perfusion, we were unable to replicate that effect *in vivo* in mice through systemic administration (data not shown)^35–41^. Future work will include both refinement of this method to induce proteinuria using less toxic substrates and enhancement of the quantity and segment of tubule cells transduced in this initial study. For the present, the gene therapy research community has a new tool for studying AAV-mediated transgene overexpression or re-expression in renal tubule epithelial cells *in vivo*. These results represent an overall important steppingstone for genetically treating tubulopathies.

## Author Contributions

J.D.R. contributed to the initial conception of the study, design of all experiments, execution of all experiments, data analysis, and drafting of the manuscript; K.A. contributed to design, execution, and analysis of flow cytometry experiments and intellectual composition of the manuscript; E.B.M. contributed to production of adeno-associated virus vectors; M.J.H. contributed to viral vector administration to animals; P.C.H. contributed to intellectual composition of the manuscript; V.E.T. contributed to the initial conception of the study, design of the induced proteinuria experiments, and intellectual composition of the manuscript; M.A.B. contributed to the initial conception of the study, design of experiments, and intellectual composition of the manuscript.

## Acknowledgements

We would like to thank Dr. Michael F. Romero for his valuable scientific feedback regarding the data presented in this paper. We would also like to thank the Microscopy and Cell Analysis Core facility at Mayo Clinic Rochester for their assistance in confocal microscopy.

## Author Disclosure Statement

The authors declare they do not have any competing or financial interests.

## Funding

This work was supported by the Department of Molecular Medicine at Mayo Clinic (J.D.R.), National Institute of Diabetes Digestive and Kidney Diseases Grant Number 1F31DK123858-01 (J.D.R.), the Department of Laboratory Medicine and Pathology at Mayo Clinic (M.A.B.) And Mayo Clinic Robert M. and Billie Kelley Pirnie Translational Polycystic Kidney Disease Center (DK090728).

## Data Availability

The datasets generated during and/or analyzed during the current study are available from the corresponding author on reasonable request.

## Supplemental Material Table of Contents

| Supplemental Fig. Number | Title |
| --- | --- |
| 1 | Example of proteinuria dipsticks used to assess induced proteinuria in mice. |
| 2 | Administration of LPS to mice did not affect liver transduction by AAV but did result in renal medullar transduction across several serotypes of AAV. |
| 3 | Evidence of toxicity associated with combined LPS and AAV administration. |
| 4 | Representative flow cytometry plots for mice administered scAAV8-Cre. |
| 5 | Representative flow cytometry plots for mice administered scAAVrh10-Cre. |
| 6 | Increased kidney transduction after administration of LPS and Ad5-Cre is negatively correlated with liver transduction. |
| 7 | Comparison of liver transduction across various vectors and doses. |

**Supplemental Fig. 1.**
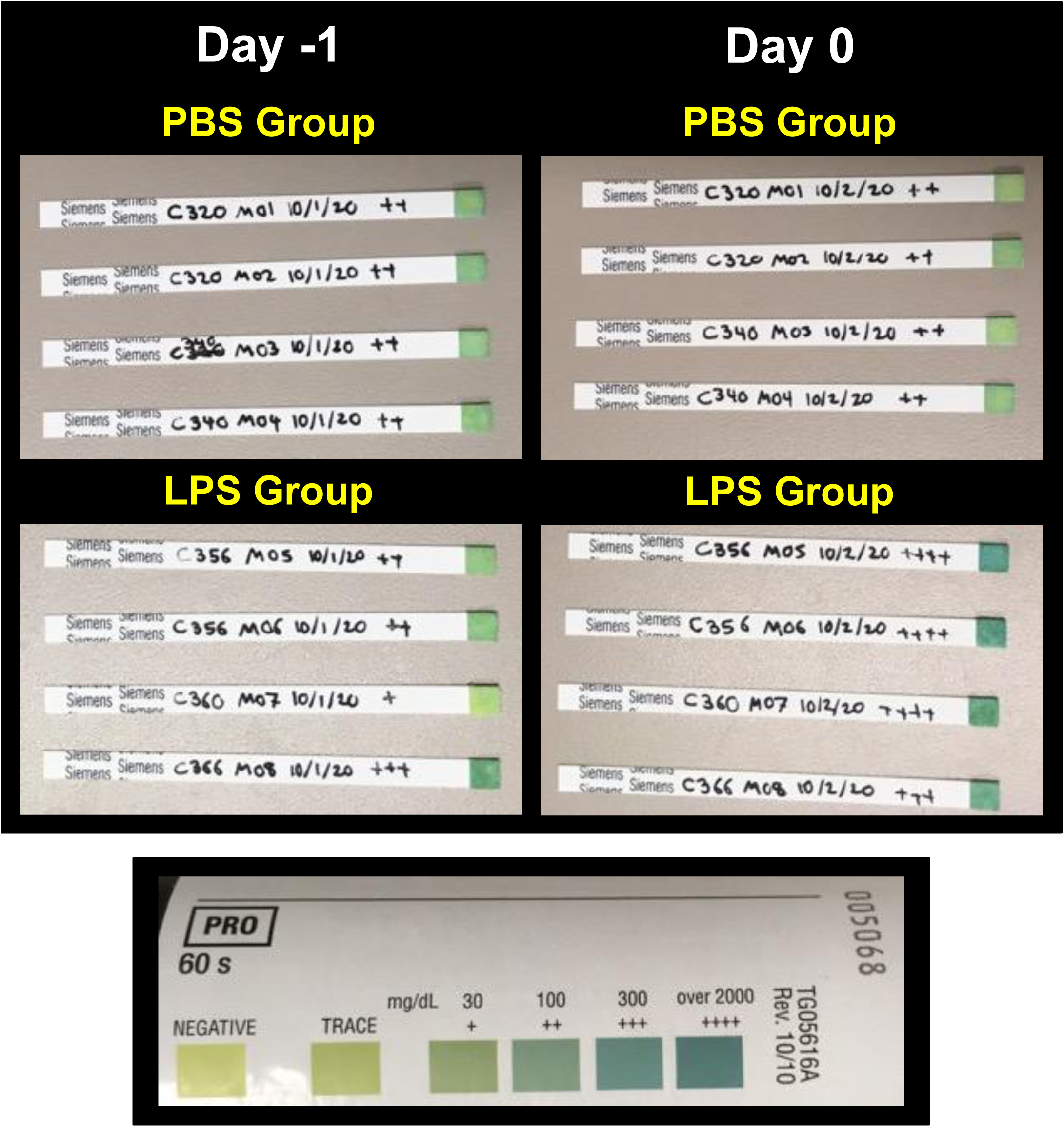

**Supplemental Fig. 1.**
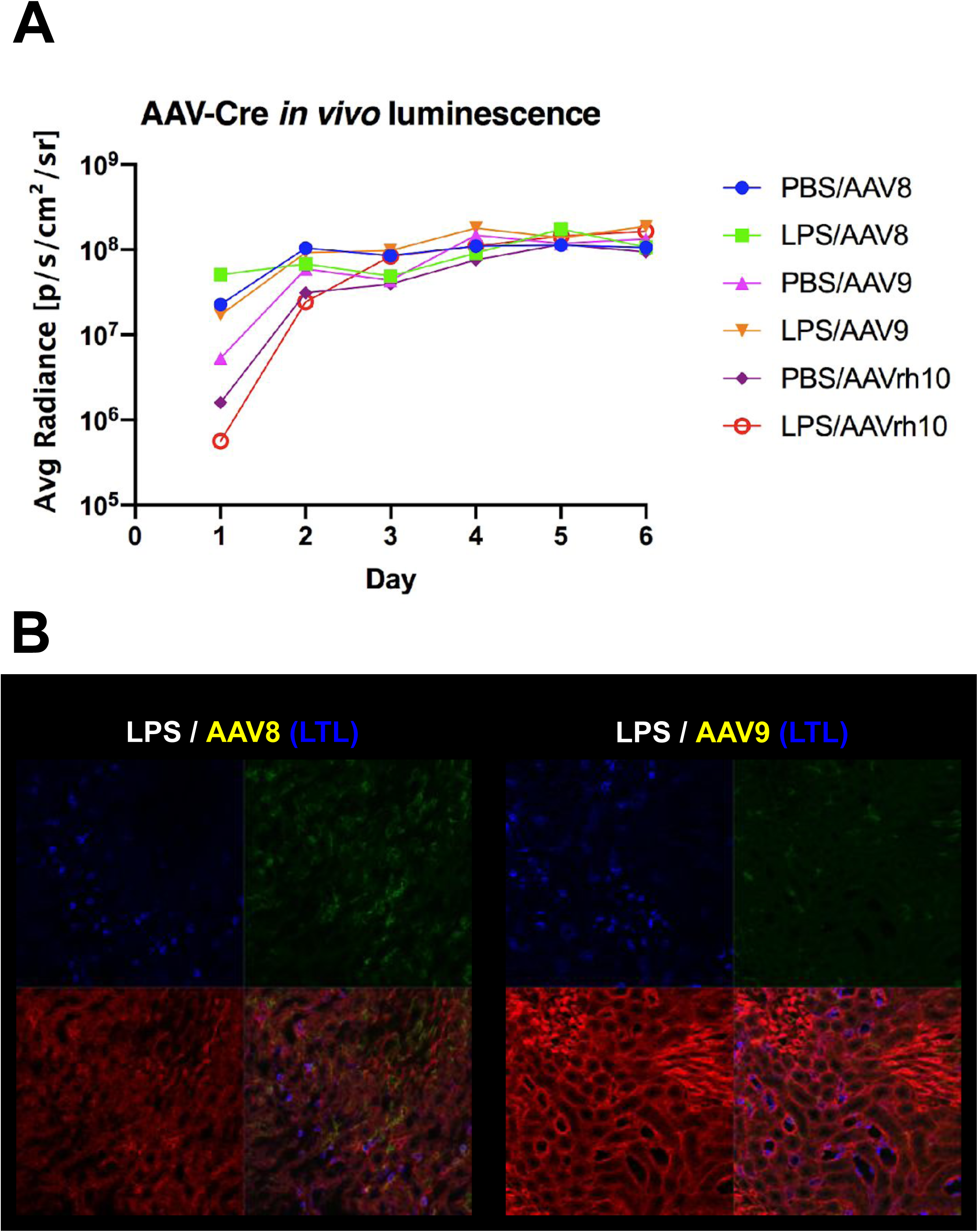

**Supplemental Fig. 1.**
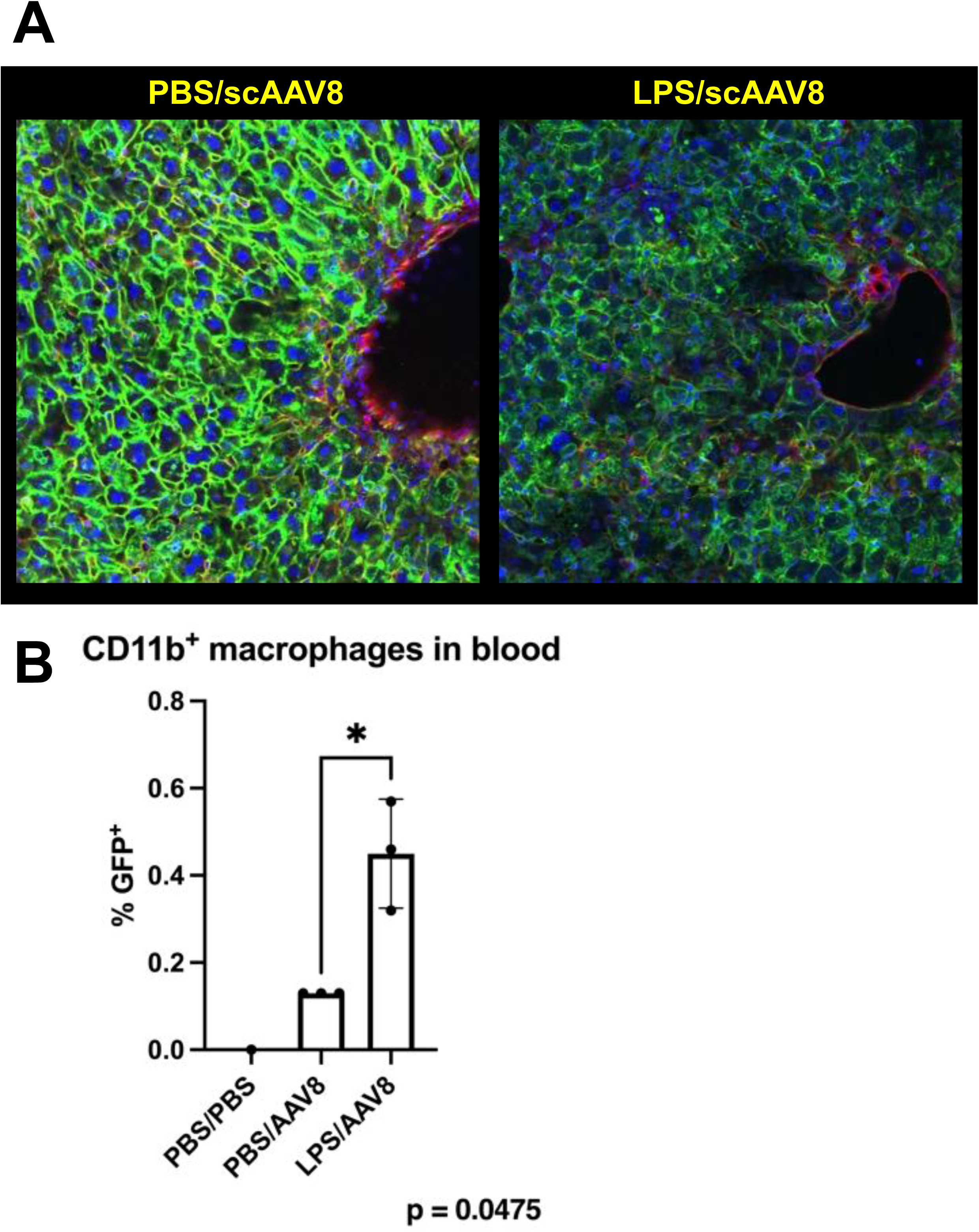

**Supplemental Fig. 1.**
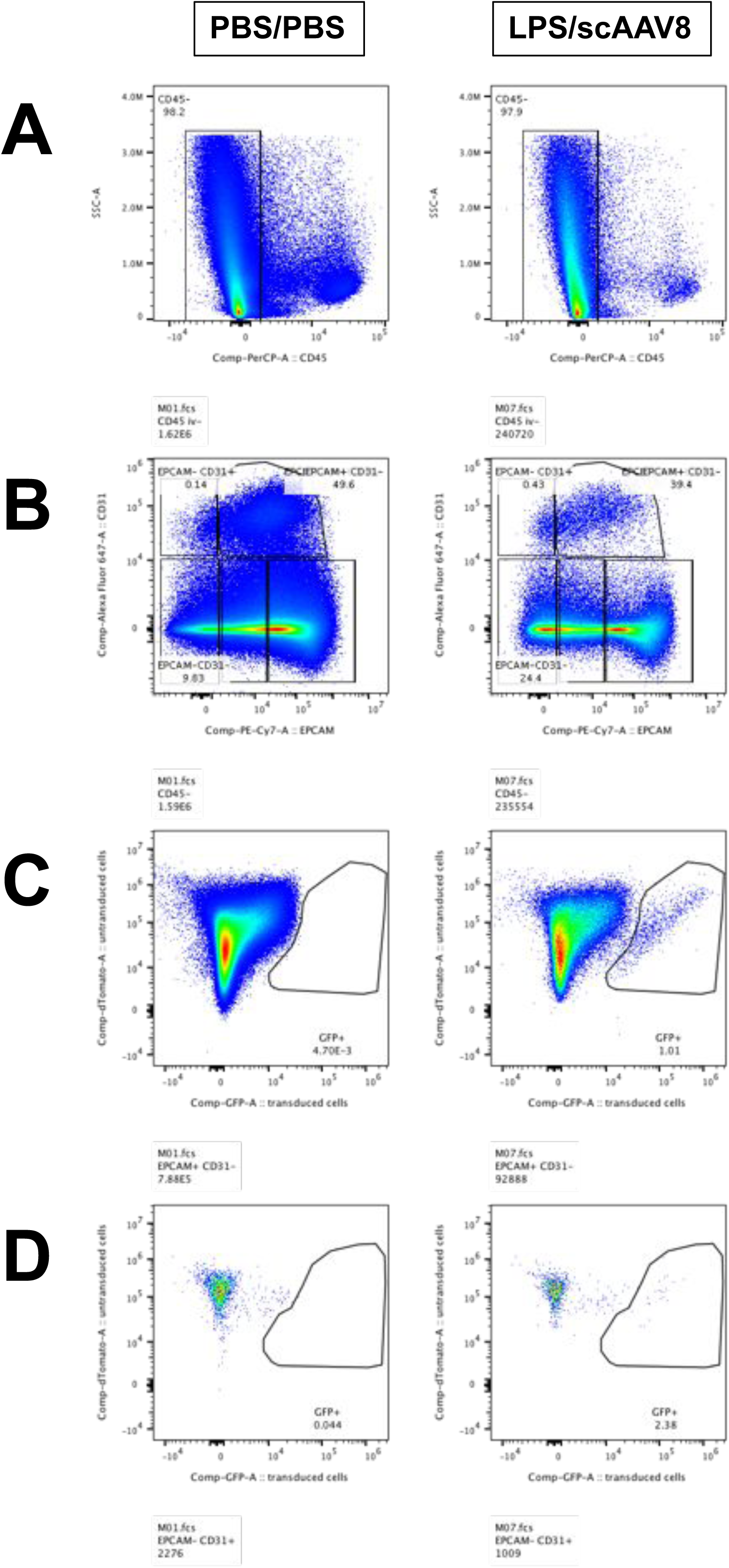

**Supplemental Fig. 1.**
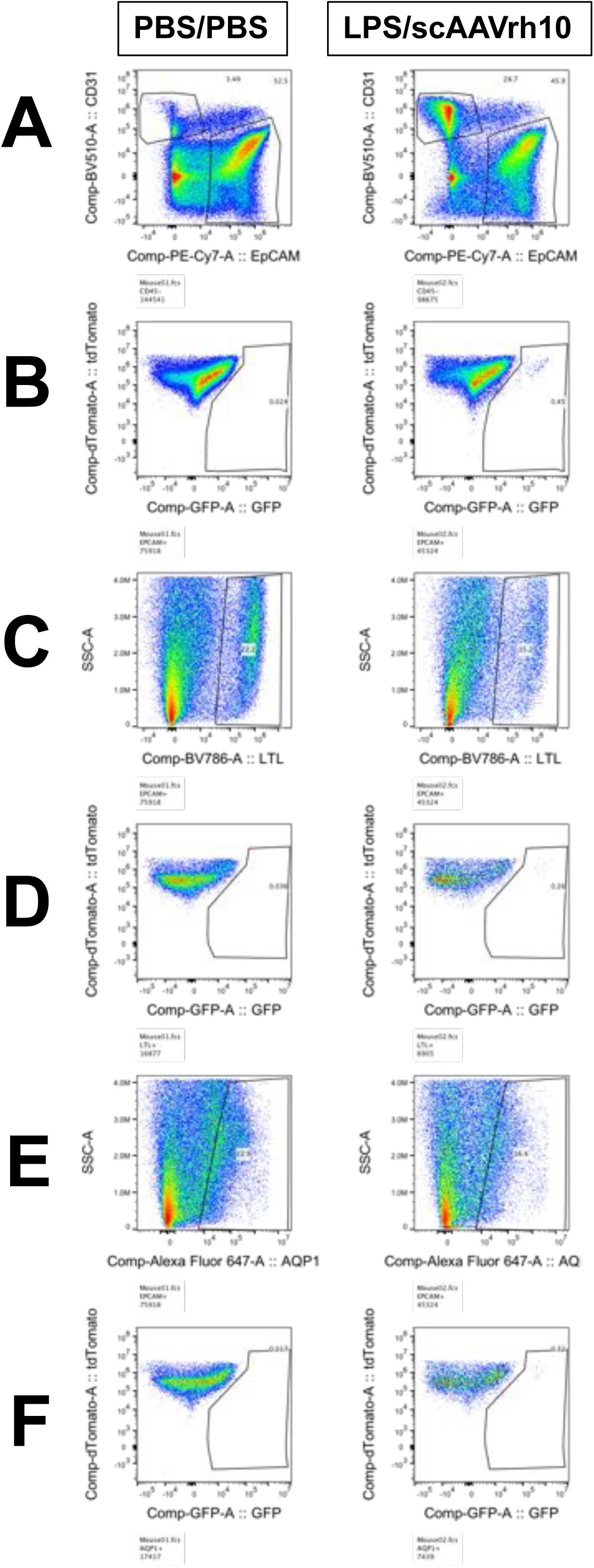

**Supplemental Fig. 1.**
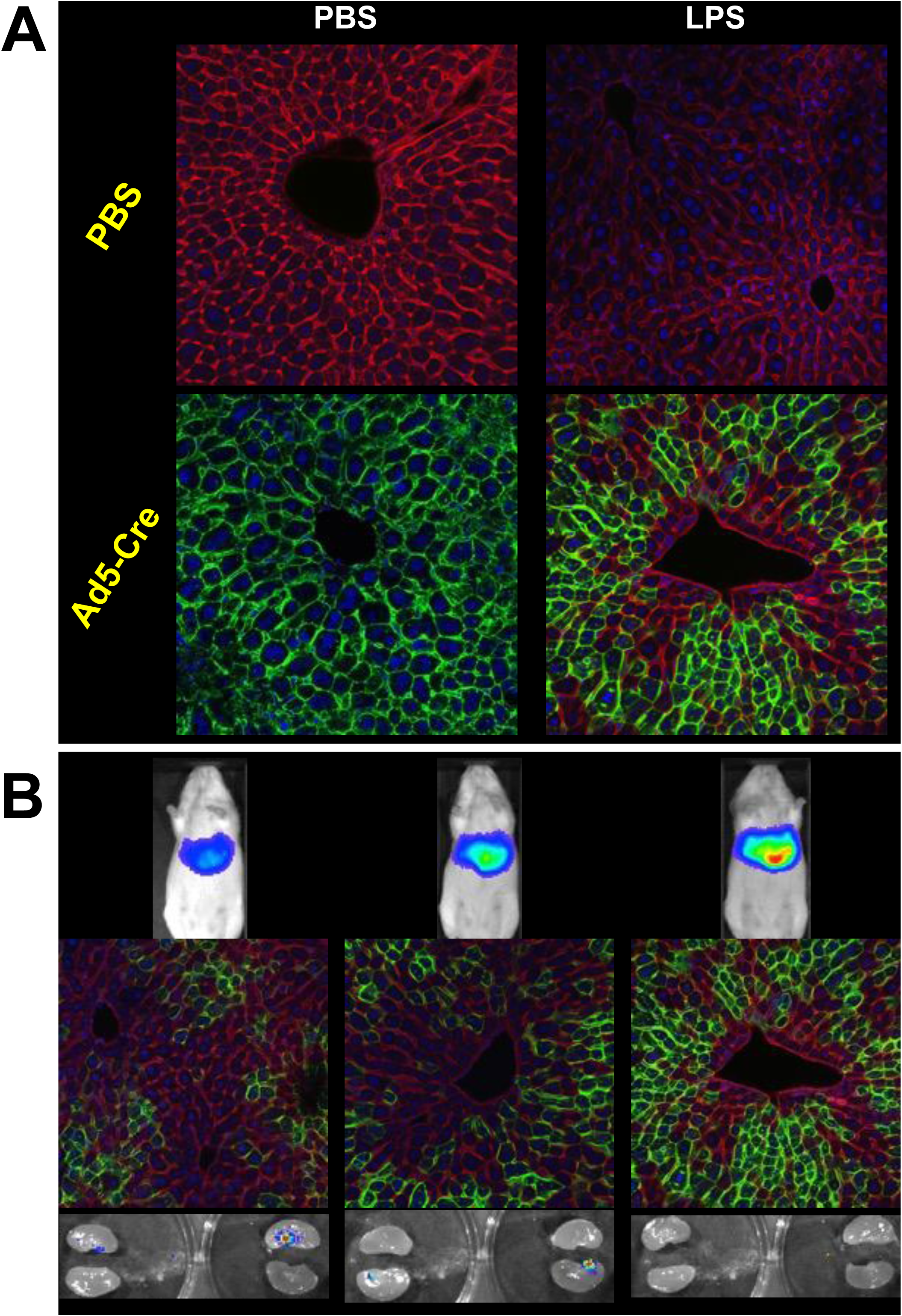

**Supplemental Fig. 1.**
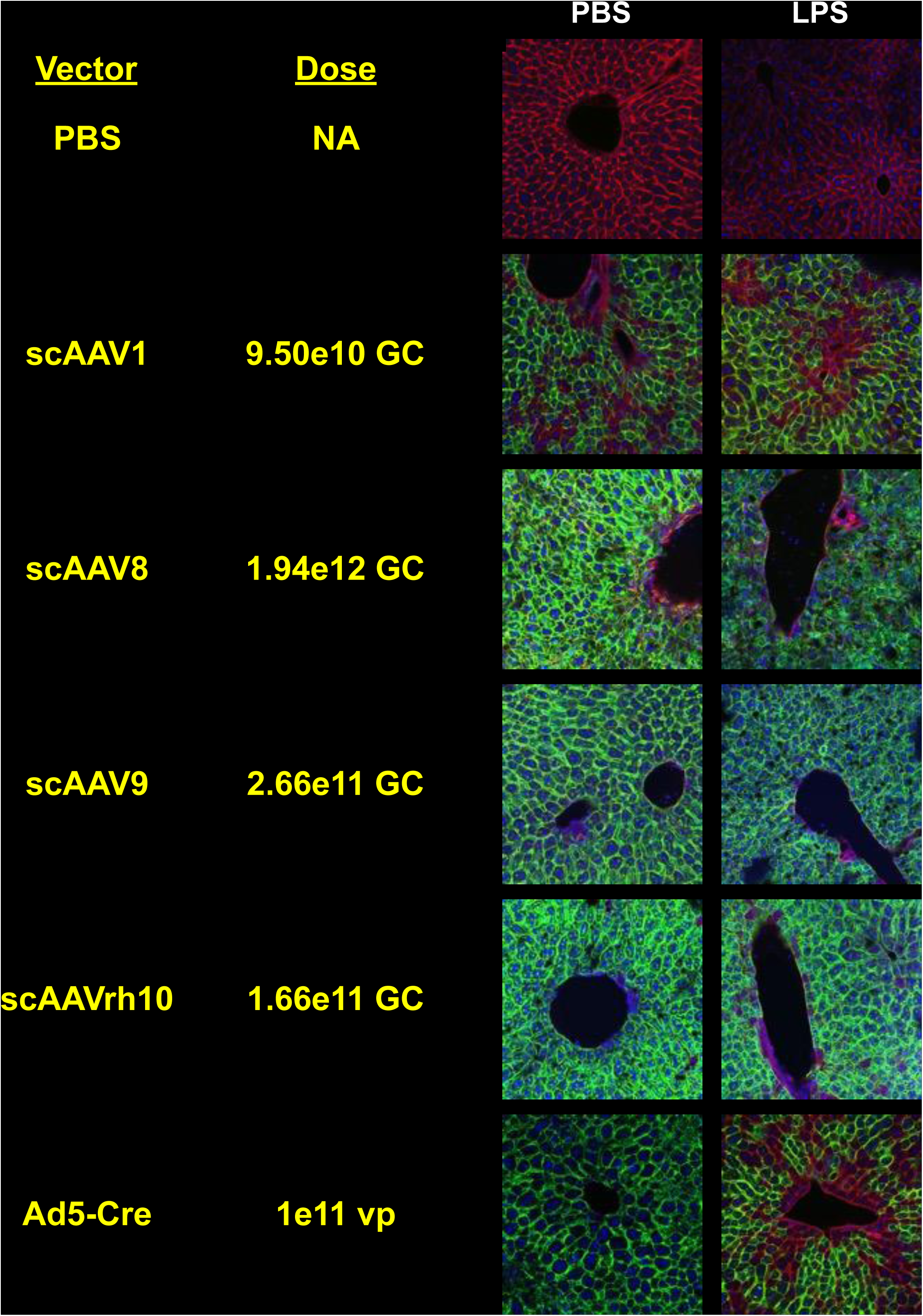

## Notes

### Competing Interest Statement

The authors have declared no competing interest.

### Summary of Updates

A review of the PDF shows that the figures were duplicated. A revision will have to be submitted to correct this. The PDF will be proofed after conversion before approving the file.

